# High-speed large-scale 4D activities mapping of moving *C. elegans* by deep-learning-enabled light-field microscopy on a chip

**DOI:** 10.1101/2021.01.19.427254

**Authors:** Tingting Zhu, Lanxin Zhu, Yi Li, Xiaopeng Chen, Mingyang He, Guo Li, Hao Zhang, Shangbang Gao, Peng Fei

**Affiliations:** School of Optical and Electronic Information-Wuhan National Laboratory for Optoelectronics, Huazhong University of Science and Technology, Wuhan, 430074, China; College of Life Science and Technology, Huazhong University of Science and Technology, Wuhan, 430074, China

## Abstract

We report a novel fusion of microfluidics and light-field microscopy, to achieve high-speed 4D (space + time) imaging of moving *C. elegans* on a chip. Our approach combines automatic chip-based worm loading / compartmentalization / flushing / reloading with instantaneous deep-learning light-field imaging of moving worm. Taken together, we realized *intoto* image-based screening of wild-type and uncoordinated-type worms at a volume rate of 33 Hz, with sustained observation of 1 minute per worm, and overall throughput of 42 worms per hour. With quickly yielding over 80000 image volumes that four-dimensionally visualize the dynamics of all the worms, we can quantitatively analyse their behaviours as well as the neural activities, and correlate the phenotypes with the neuron functions. The different types of worms can be readily identified as a result of the high-throughput activity mapping. Our approach shows great potential for various lab-on-a-chip biological studies, such as embryo sorting and cell growth assays.

## Introduction

Microfluidics technology developed over the past few decades has greatly impacted biomedical research, therapeutics, and diagnostics. Various applications, such as small compound screening and gene expression studies, greatly benefit from increased throughput and sensitivity provided by microfluidics^1–6^. Generally, a large-scale microfluidic assay also requires massive measurement of samples, for example, fluorescence-activated cell sorting (FACS), through which followed-up analyses, such as counting and phenotyping, are further enabled^7–11^. Light microscopy techniques, especially well-established epifluorescence microscopes, have been nowadays widely used for various on-chip assays containing biological specimens from single cell to multi-cellular organisms. However, as many millisecond transient cellular processes occur in three-dimensional (3D) tissues and across long time scales, a recurring challenge for on-chip measurement is the quest to extract ever more spatiotemporal information from targets at sufficiently high throughput. While the classic microscopy techniques, including epifluorescence and plane illumination methods, can image live samples in three dimensions at high spatial resolution^12–16^, they require recording a number of two-dimensional (2D) images to create a 3D volume, and the temporal resolution is compromised by the extended acquisition time of camera. Recently, light-field microscopy (LFM) has become the technique of choice for instantaneous volumetric imaging^17–24^. It permits the acquisition of transient 3D signals via post-processing of the light-field information recorded by a single 2D camera snapshot. Because LFM provides high-speed volumetric imaging limited only by the camera frame rate, it has delivered promising results for various applications, such as the recording of neuronal activities^18–21^ and visualization of cardiac dynamics in model organisms^22^. While LFM is suited for the volumetric imaging of dynamic processes, its previous applications are restricted to the recording of single live sample, which is still insufficient for conducting large-scale biological assays that requires measuring a number of samples in a short time.

Here we present a hybrid approach that integrates light-field microscopy with microfluidic chip to achieve high-speed, high-throughput activity mapping of live *C. elegans* at large scale. A multi-layer microfluidic chip with control valves is designed to rapidly load worm samples, sequentially compartmentalize them one by one into a chamber for sustained light-field recording, and flush them one by one to the outlet for collection^25–27^. A customized light-field microscope combined with a deep-learning-based reconstruction algorithm further enables the real-time capture and 3D restoration of the behaviours and neural activities of each compartmentalized moving worm. We demonstrated that an assay mixed with wild-type (WT) and uncoordinated-type (Unc) *C. elegans* are *intoto* imaged and quantitatively analysed using this approach at a throughput of 42 samples per hour.

## Results and discussion

### Experimental Setup

We built a platform for high-throughput automatic *C. elegans* trapping, imaging and analysis (Fig. 1). A valve-based microfluidic chip was designed to manipulate the *C. elegans* via adapting its channel geometries to the size and shape of the worms. A LFM system was specifically designed to record the activities of moving *C. elegans* inside the chip at high speed. In the LFM setup, a microlens array (MLA) was placed on the plane conjugating to microscope’s native image plane to allow the light-field recording (Fig. S1), by which a 3D volume could be reconstructed through a single 2D capture. During experiment, a number of live *C. elegans* larva (L4 stage) were loaded into the chip from the reservoir, and sequentially trapped into the chip’s imaging chamber (~600 × 600 × 50 μm) for 1 minute (Fig. S2), during which each worm was crawling inside the chamber while simultaneously imaged by the LFM system at a high acquisition rate of 33 frames per second (fps, up to 100 fps). The fluorescencely-labelled neural activities (GCaMP *Ca*^*2+*^ indicator) of the behaving worm were thus recorded instantaneously. After the on-chip tasks, the acquired raw light-field videos of all the worms were computationally reconstructed by a light-field recovery program to yield the 4D visualizations (in 3D space + 1D time) of the worm activities. Given the fact that classic light-field deconvolution (LFD) algorithms suffered from limited reconstruction quality and low computation speed^18^, here we used a deep neural network (DNN, U-Net^28^) to quickly transform the raw 2D light-field images into a sequence of 3D volumes with quality notably better than light-field deconvolution method^29^. Such a DNN model was iteratively trained on the high-resolution 3D reference images of the static worms and their corresponding light-field projections (LFP) which were projected from the 3D reference images based on wave optics model^24^ (Fig. 1b), learning how to transform the views of input light-field measurements into depth images of a 3D output through correlating them with the channels of the DNN. Finally, Such a well-trained view-channel-depth (VCD) transformation model can directly infer a sequence of high-resolution 3D images from the raw light-field measurements at a high rate of ~13 volumes per second.

**Fig. 1.**
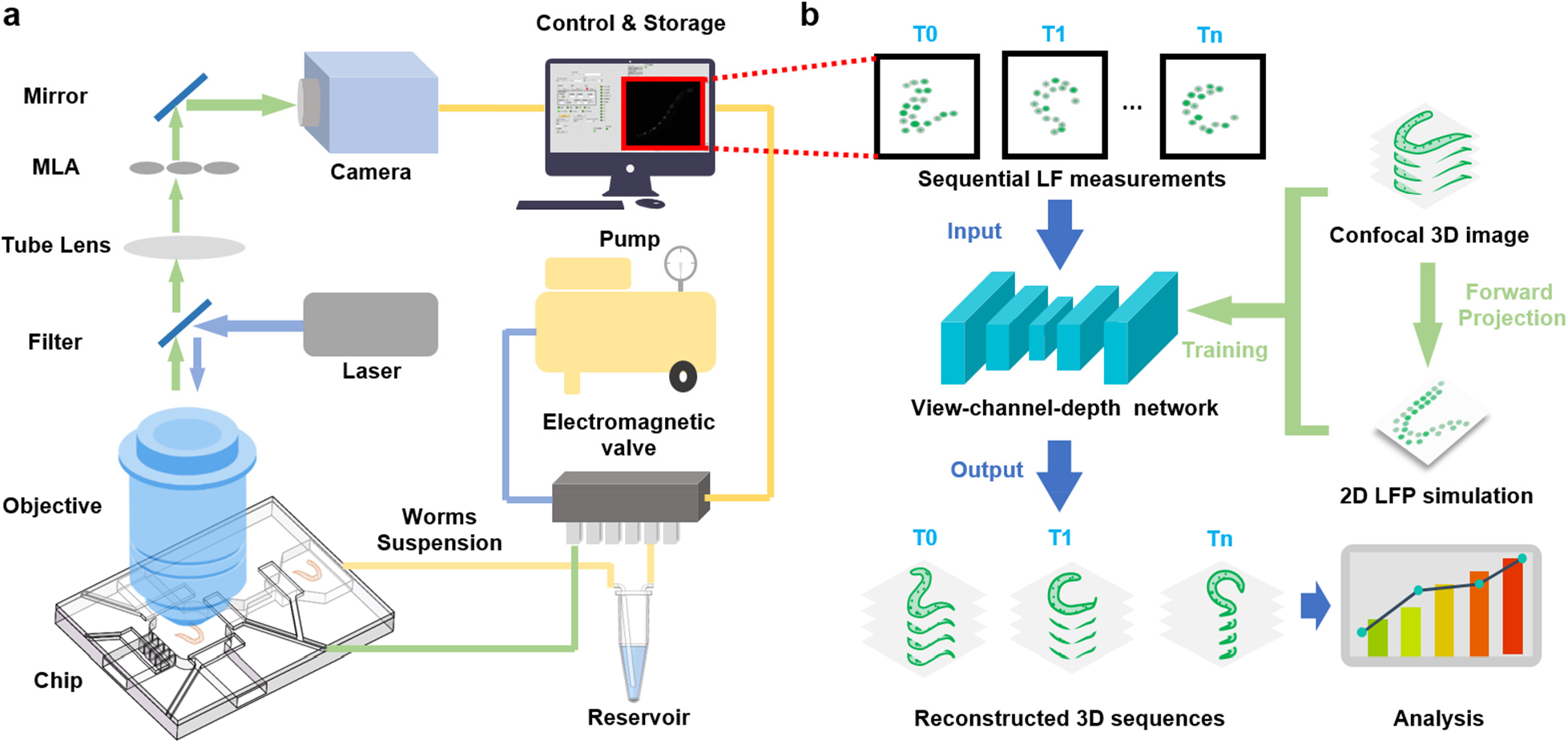
On-chip light-field microscopy platform for high-throughput *C. elegans* trapping, 3D imaging and quantitative analysis. (a) Schematic of the worm manipulation and imaging. The two-layer microfluidic chip is operated with electromagnetic valves to sequentially transport the *C. e/egans* larva from a reservoir to an imaging chamber (600 × 600 × 50 μm), the size of which is matched with objective's field of view (Olympus 20x/0.5 water dipping objective). During the 1-minute stay in the chamber, each mocving C. elegans is imaged by a self-built light-field microscopy system, and then flushed away when the observation finishes. A new worm will be loaded inside the chamber to repeat this process. The valves and the camera are synchronized by a LabVIEW program to allow the implementation of whole process in good order. (b) Schematic of light-field reconstruction and image-based quantitative analysis. Sequential light-field (LF) images are processed by a view-channel-depth (VCD) network, which has been trained by confocal 3D images paired with 2D light-field projection (LFP) simulation. Reconstructed 3D sequences are subsequently used for further analysis.

### Microfluidic chip design

The microfluidic chip worked on three modes as: worm loading, trapping and flushing, which could be switched automatically by LabVIEW program. The flow layer of the chip contained an imaging chamber with 4 fluid channels, *C. elegans* loading inlet, flushing inlet, flushing outlet, and water outlet, connected from four directions (Fig. 2a, b). Under the control of micro valves at control layer, the worms were sequentially loaded into the chamber and trapped inside chip’s imaging chamber for light-field imaging of their neural activities in an automatic way (Fig. 2c, Supplementary Video 1,2). A fence-like structure was designed in port 2 (Fig. 2a) to prevent the worms from escaping. To allow the loading/washout of worms in good sequential order, we narrowed down the width of fluid channel from ~600 μm at the main part into ~50 μm at connecting part, a size approximately the width of the worm body, thus allowing only single worm to be pushed into the chamber at a time. The 600 × 600 μm sample chamber with size matching the field-of-view (FOV) of our LFM (20×/0.5w objective and Hamamatsu Flash 4.0 v2 camera) was closed by valves to trap the *C. elegans* inside when imaging continued, and opened by valves to wash out the worm after imaging finished. A LabVIEW program automatically switched the chip among these three modes through judging whether there was a worm in the chamber and how long it had stayed there. The workflow of LabVIEW program was also shown in Fig. S3. In our demonstration, totally 42 moving worms mixed with WT worms and Unc mutants containing motion deficiency were volumetrically imaged (2000 image volumes for each worm) and analysed using our microfluidics-based LFM platform in less than 1 hour.

**Fig. 2.**
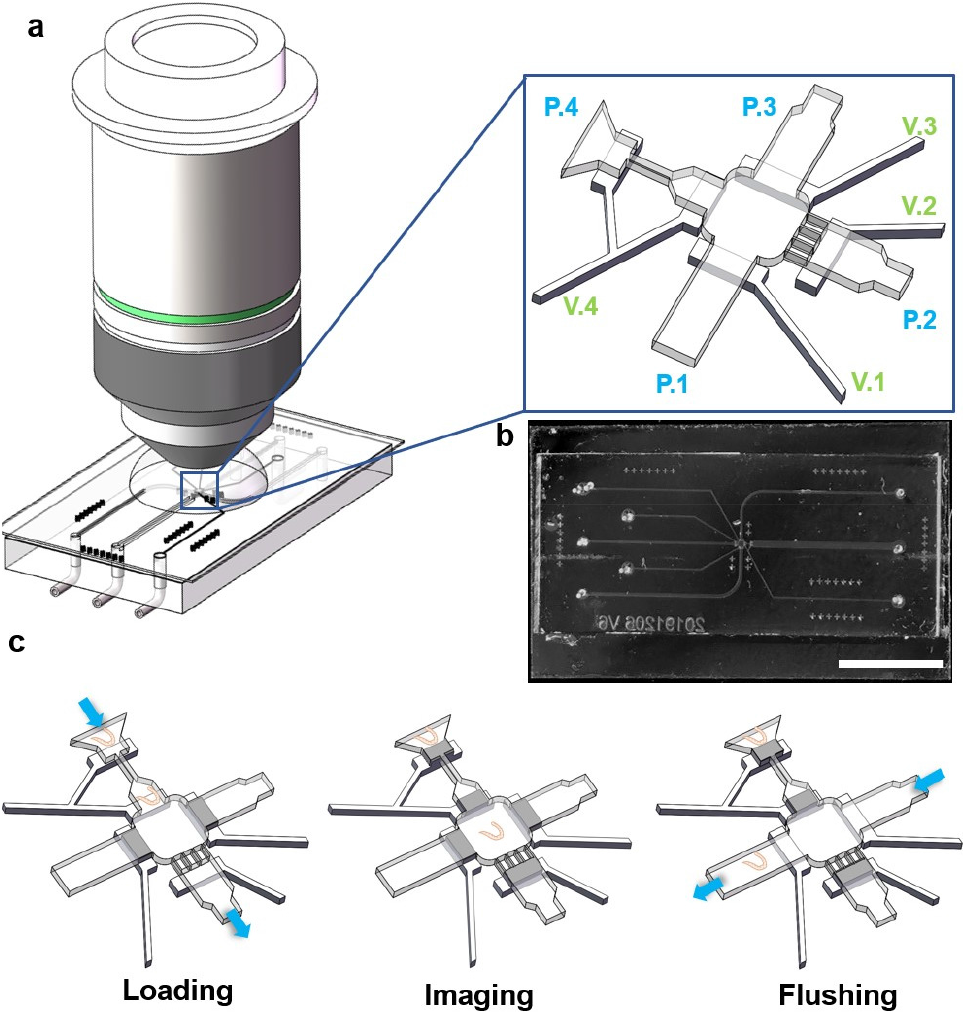
Microfluidic chip design and valves control process. (a) Design for sequentially loading the worms inside the micro chamber for live light-field imaging. The chamber is connected with four ports P.1-P.4 (magnified view), which can be opened or closed by four micro valves V.1-V.4, respectively. (b) Photograph of a fabricated chip. Scale bar 10 mm. (c) Operating procedure for loading, imaging and flushing the *C.elegans* larva. At loading status, P.1 and P.3 are closed and *C. e/egans* are loaded from P.4. P.2 is also opened as buffer output. At imaging status: Once a worm is loaded into the chamber, all the four ports are closed to trap the worm for one minute. Meanwhile, the camera starts to record the light-field images at video frame rate. The chip is switched into flushing status after imaging. P.1 and P.3 are opened and the current worm is flushed outside the chamber by buffer from P.3 to P.1. P.1, P.3 are then closed, and P.2, P.4 are opened again for another cycle of loading a new worm.

### Light-field measurement and deep-learning reconstruction

We used a 20×/0.5w objective to image the motor neuron activities of L4-stage *C. elegans* larva (strain ZM9128 *hpIs595*[*Pacr-2(s)*::GCaMP6(f)::*wCherry*] and SGA197 *unc-13(s69)*; *hpIs595*[*Pacr-2(s)*::GCaMP6(f)::*wCherry*]) with 5-ms exposure time and 33-Hz acquisition rate, yielding 2000 light-fields in a 1-minute observation for each worm. The deep-learning-enabled light-field recovery then provided high-quality and quantitatively-accurate 3D reconstructions to *intoto* visualize the neuron calcium signalling of whole *C. elegans* at single-cell resolution during fast body movement (Fig. S4, Supplementary Video 3). The VCD network model developed for high-performance light-field recovery is based on well-established U-(Fig. 3), which contains subpixel convolutional layers, different convolutional layers, and deconvolution layers. To pair the net^28^ views of the light-field projections with the depths of the high-resolution 3D target, we designed several subpixel convolutional layers to interpolate the extracted views into the size of 3D target images before the U-Net training. There are four subpixel convolutional layers with each up-scaling the image by a factor 2. We adopted 5 convolutional layers for feature extraction and 5 deconvolution layers for image reconstruction. Compared to either wide-field microscopy which can only record neuron signals on a single focal plane, or confocal microscopy which can perform 3D scanning but at low speed insufficient for instantaneous imaging, our method volumetrically records all the neurons of moving *C. elegans* while noted that conventional confocal microscope typically works at mega voxels per second throughput, our method can yield ~1 Giga voxels per second, over two-order-of-magnitude faster than commonly-used 3D imaging method.

**Fig. 3.**
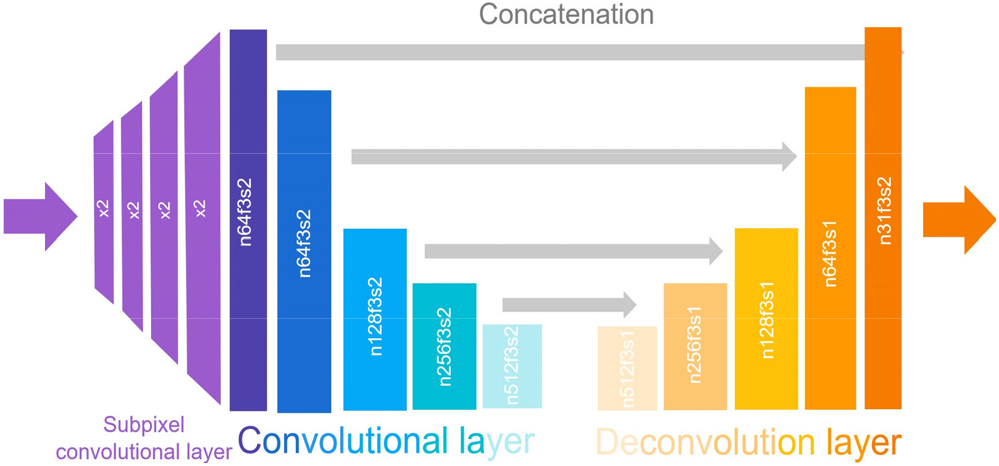
Network architecture of the VCD light-field reconstruction model. The model contains 4 subpixel convolutional layers, 5 convolutional layers and 5 deconvolution layers. The hyper-parameters of each convolutional layer and deconvolution layer were indicated by n, f, and s, which represented the number of output feature maps, the convolutional kernel size and the strides, respectively.

### Locomotion patterns and neural activities of moving WT and Unc mutant worms

The deep-learning-enabled LFM in conjunction with chip-based sample manipulation enables sustained recording of *C. elegans*’ instantaneous positions in three dimensions (Supplementary Video 4), as well as high-throughput on-chip screening of a large number of worms. As shown in Fig. 4, the visualized trajectories of 6 Unc mutant worms (7-12) are significantly different from those of 6 WT worms (1-6), in term of their larger body curvatures, lower velocities, and shorter locomotion distance. Besides the spatial distribution, to further investigate the neural activities reflected by the fluctuation of *Ca*^*2+*^ signals, the intensity of light-field reconstruction needs to be quantitatively accurate as well. We validated this through tracing the intensity fluctuations of reconstructed signals and compared them with the ground truths. The high similarity shown between the signal fluctuation curves has verified the intensity accuracy of reconstructed signals and their ability to indicate the neural activities correctly (Fig. S5). Then, for both WT and Unc mutant worms, we specifically identified their A- and B-motor neurons been associated with motor-program selection (Fig. 5a, Fig. S6), and mapped their calcium activities over time (Fig. 5b). Furthermore, through applying an automatic segmentation of the worm body contours based on the location and that have quantitatively analysed the changes of curvature and velocity related to worm locomotion and behaviour (Fig. 5c, d). It is quite easy to tell the significant differences between WT and mutant worms in term of both their locomotion and neural activities, from which we can readily sort the WT worms with normal motion ability and Unc worms with motion deficiency. Finally, we identified 33 WT worms and 9 Unc worms from totally 42 inputs, and *intoto* mapped out their behaviours and neural activities.

**Fig. 4.**
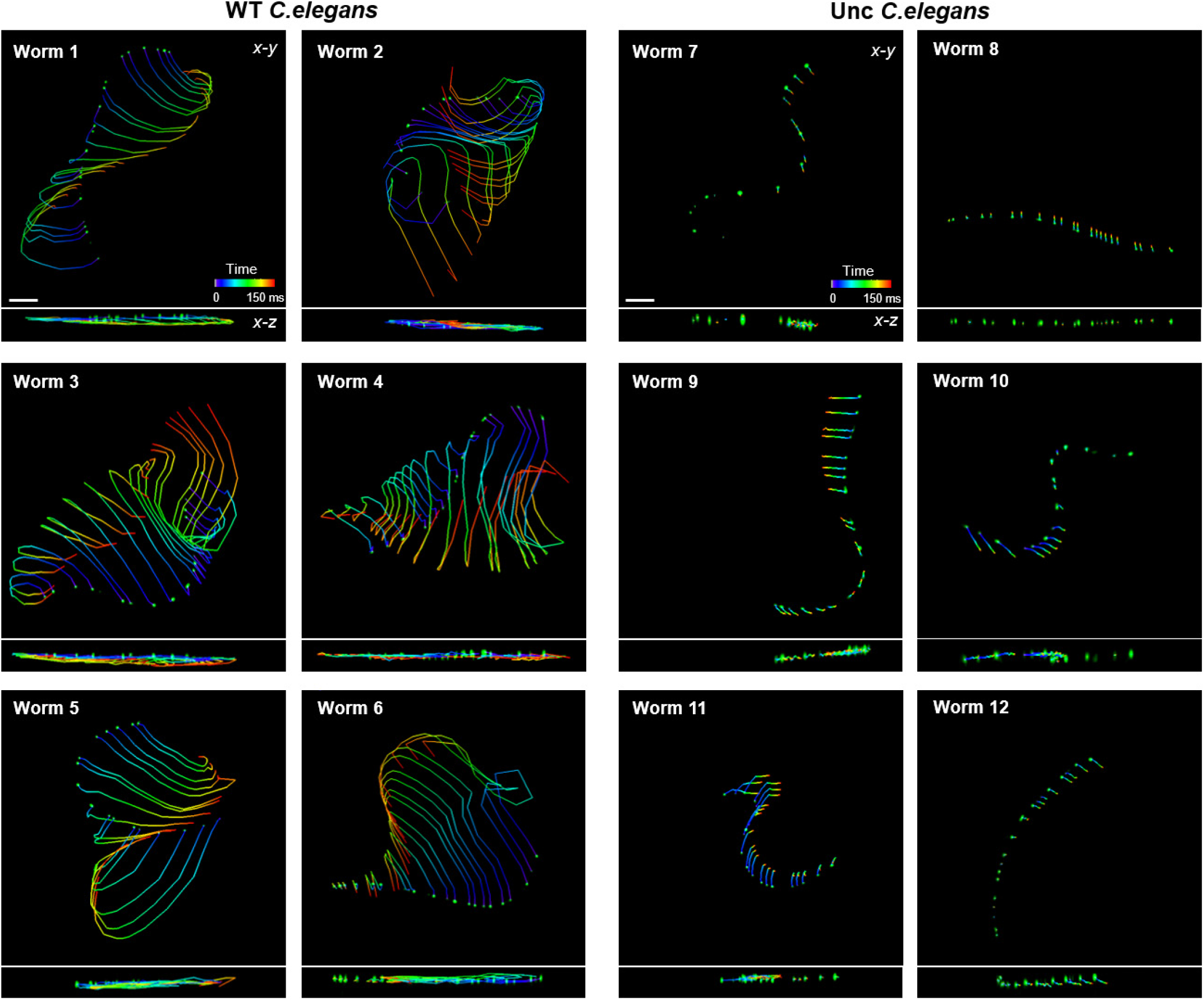
3D traces of the motor neruons in moving WT (strain ZM9128 *hpls595[Pacr* 2(sJ::GCaMP6(f)::wCherry]) and Unc mutant (strain SGA197 *unc-13(s69}; hpls595[Pacr-2(s)::GCaMP6(f)::wCherry]) C. elegans* larva (L4 stage). The tracing was performed based on the 4D imaging of the worm cocktail (mixed with 33 WT worms and 9 Unc worms) using our microfluidics-based LFM platform. Six worms were selected from each type to visualize their 3D trajectory and velocity in 150 ms. It's obvious that WT *C. e/egans* are much more active than Unc *C. e/egans.* The colormap indicates the time-varying positions of the neurons. Scale bars, 30 μm.

**Fig. 5.**
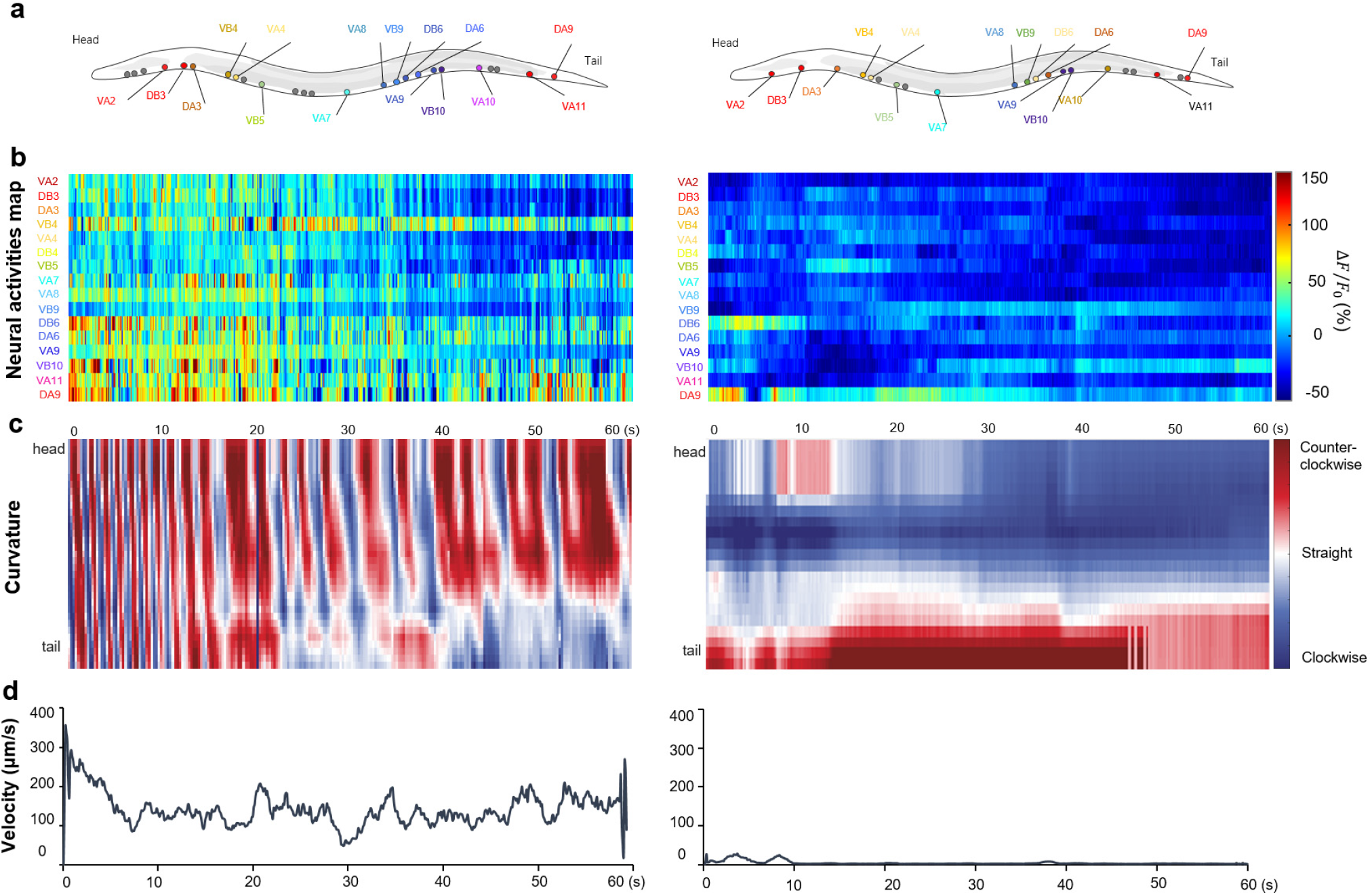
Quantitative analysis of the motor neuron activities and corresponding locomotion behavior of the WT and Unc mutant C. *elegans.* (a) Soma position of the motor neurons in WT (left) and Unc (right) *C.elegans* larva, respectively. (b) Activity of the 16 motor neurons identified in (a). Each row shows a time-series heat map indicating the percent fluorescent fluorescence changes {8F / Fa) of an individual neuron. (c) Representative kymographs of bending curvature along the body of the moving worms. (d) Velocity plot of the moving WT and Unc worms during 1 minute.

## Conclusions

Our microfluidics-based LFM approach is the first proof-of-concept that shows the high-throughput live sample manipulation in conjunction with instantaneous 4-D imaging of dynamic processes, both of which could be beneficial to large-scale assay applications. From the perspective of imaging, the chip-based automatic sample loading and manipulation greatly facilitate the implementation of LFM and improves its throughput. On the other hand, the introduction of LFM also empowers the lab-on-a-chip system with advanced imaging capability, for volumetric imaging-based *intoto* screening and analysis. Thus, the synergistic combination of LFM measurement and microfluidics could maximize the efficiency of various live imaging assays. Using this approach, we successfully demonstrated high-speed 4D screening on *C. elegans* managed with different genotypes and phenotypes. The dynamic worms’ locomotion, neural activities are volumetrically recorded and quantitatively correlated to sort the worms according to the assessment on their personalized behaviours. Our approach thus provides a paradigm shift which enables *intoto* activity mapping of dynamic specimens, a challenge unmet by other alternative approaches. Considering the highly flexible design of microfluidic chip for various purposes, this approach shows good potentials for benefiting a variety of large-scale, lab-on-chip assays.

## Acknowledgements

This research has received the funding support from China 1000 Youth Talent Plan (P.F. and S.G.), the National Natural Science Foundation of China (21874052, 21927802 and 31871069), and the innovation fund (L.Z., 2020JYCXJJ076). The authors would like to thank Mei Zhen, Honghao Ma for their helps on the development of *C.elegans* strains and the chip fabrication.

## Notes

### Competing Interest Statement

The authors have declared no competing interest.

