## Supplementary Materials for "High-speed large-scale 4D activities mapping of moving *C. elegans* by deep-learning-enabled light-field microscopy on a chip"

### Supplementary Information

#### Methods

##### Microfluidic Device Fabrication

We prepared a three-layer microfluidic device using soft lithography. We first made silicon mold masters using conventional photolithography, forming a 60-μm-thickness photoresist pattern (Microchem, SU8 2050) for fluid layer, and a 50-μm-thickness photoresist pattern (Microchem, SU8 2025) for control layer. Before chip fabrication, silicon mold masters were treated with trimethylchlorosilane for hydrophobicity. The fluid layer of microfluidic chip was fabricated by casting polydimethylsiloxane (PDMS) (Sylgard 184 A and B, 1:5 by weight, Dow Corning) on fluid layer master. The thickness of fluid layer was ~4 mm. The control layer was fabricated by spinning coating a thin layer of PDMS (~100 μm, 1:20 by weight) on control layer master. The spinning speed was first 500 r/min for 12 s and then 1600 r/min for 40 s. Both layers were then partially cured at 80 ℃ for 20 min. The fluid layer was then peeled off, with inlet and outlet holes being punched using a 0.6-mm dermal punch. Then the fluid layer was aligned with the control layer and thermally bond together at 80 ℃ for 40 min. After baking, the control layer bonded with fluid layer was peeled off from the mold master with the inlet and outlet holes at control layer were punched using a 0.6-mm dermal punch. Finally, the PDMS chip containing fluid and control layers was bonded to a cover glass that was prior coated with a thin layer of PDMS (~80 μm, 1:10 by weight, 3500 r/min spin coating for 60 s). The fabricated chip was cured overnight at 80℃ before use.

##### LabVIEW Control Software

To achieve light-field imaging of *C. elegans* at high throughput, we designed a LabVIEW control program to realize worm loading, imaging and flushing automatically. The schematic of LabVIEW program was shown in Fig. S3. The program could control the camera and solenoid valves and switch the system among three stages of the cycle from flushing the old worm, to loading a new worm, and to imaging the new worm. The automatic switch was activated based on the identification of worm status in the chamber. We used the maximum intensity of live images to make real-time judgement of whether there was a worm inside the chamber. The imaging parameters were pre-set and frames were automatically acquired at imaging stage, after which the current worm would be flushed out and a new worm would be pumped in automatically.

##### Experimental Setups

The microfluidics-based LFM system consisted of three major parts as: light-field microscopy imaging, microfluidic chip environment and automatic fluid control by solenoid valves. The valves and camera are controlled by a LabVIEW program to realize programed *C. elegans* loading, compactization, light-field recording and flushing in a sequential mode. Before connection between solenoid valves, the chip was first cleaned using ultrasound for 30 min and baked at 80 oven to remove potential impurity in channel. Then chip was connected to solenoid valves by 0.6-mm-diameter tygon tubes. The micro valves of the chip were then filled with water by opening the solenoid valve to pump the water into control layer. Before loading *C. elegans larva*, air bubbles inside the chip were removed by closing the water outlet channels (P. 2), flushing inlets (P. 1), and outlet channels (P. 3). Then the water was pumped into the imaging chamber from the loading inlet channel (P. 4) slowly. Once the air bubbles were wiped off, the loading stage would start and the subsequent steps would be executed by LabVIEW program automatically, as elaborated in legend of Fig. 2. Video S1 further shows the working procedure of the chip.

**Strain.** The strain ZM9128 *hpIs595[Pacr-2(s)::GCaMP6(f)::wCherry]*, with calcium indicators tagged to the A- and B- class motor neurons, was used as the wild-type (WT) worm that showed normal behaviors. The uncoordinated-type (Unc) strain SGA197 *unc-13(s69); hpIs595* was generated when chemical synaptic transmission was further eliminated from the entire nervous system. All *C. elegans* were cultured on standard Nematode Growth Medium (NGM) plates seeded with OP50 and maintained at 22 °C incubators until L4 stage.

**Image acquisition.** Light-field images and wide-field images were acquired using the system shown in Fig. S1. A water immersion objective (Fluor 20×/0.5w, Nikon) were used to collect the $Ca^{2+}$ fluorescence signals from the moving *C. elegans* in the microfluidic chip. The imaging time for each worm was 1 minute. The sCMOS camera worked at 33 frames per second, correspondingly recorded 2000 light-field frames for each worm (2048 × 2048 pixels in each frame). A dichroic mirror enabled (Fig. S1) the switching between 2D light-field imaging and 3D wide-field imaging, through which we could validate the quality of light-field reconstruction by view-channel-depth (VCD) network through the comparison with wide-field result (Fig. S4). To acquire high-resolution (HR) 3D images for network training, the L4-stage anesthetized worms (QW1217 *hpIs467*, ZM9128 *hpIs595*, by 2.5 mM levamisole in M9 buffer, Sigma-Aldrich) were imaged by a confocal microscope (FV3000, Olympus) using a 40×/0.95 objective.

**VCD network reconstruction.** For VCD network training, HR 3D images of anesthetized worms acquired from confocal microscopy were first projected into the light-field projections based on wave optics model[1]. Then the HR 3D confocal data and corresponding 2D synthetic light-field projections were input into the VCD as training datasets (3180 training pairs of size 176 × 176 × (31) pixels). After iteratively minimizing the difference between the intermediate 3D outputs and the HR confocal references, the well-trained VCD-Net could infers series of 3D images from a sequence of light-field images of the *C.elegans* freely moving in the microfluidic chip. As a reference point, it took ~3.5 hours for training the model (110 epochs) using a single Nvidia 2080Ti GPU. Then it merely took ~2.5 minutes for the VCD network to reconstruct 2000 light-field images, which intoto recorded 1-minute neural activities and behavior of one worm.

**Quantitative Analysis of neural activities and locomotion behavior of moving worms.** We performed semi-automated tracking of the movement and intensity fluctuation of each individual neuron using TrackMate Fiji Plugin[2]. 5-μm circular regions of interest (ROIs) were automatically detected, and the pixelwise averages within these ROIs in each time frame were calculated as $\mathrm{F(t)}$. The averages of the $\mathrm{F(t)}$ in all the time frames were correspondingly defined as $\mathrm{F_0}$. Then the $Ca^{2+}$ fluctuation were calculated as $\Delta\mathrm{F} \ / \ \mathrm{F_0}$, where $\Delta\mathrm{F} = \mathrm{F(t)} - \mathrm{F_0}$. The curvature and velocity were measured based on the same fluorescence images. We first segmented the worm outlines by enhancing the contrast and making binary of the images. Then we calculated the curvatures and velocities of the moving worm based on the segmented worm outlines.

#### Supplementary Figures

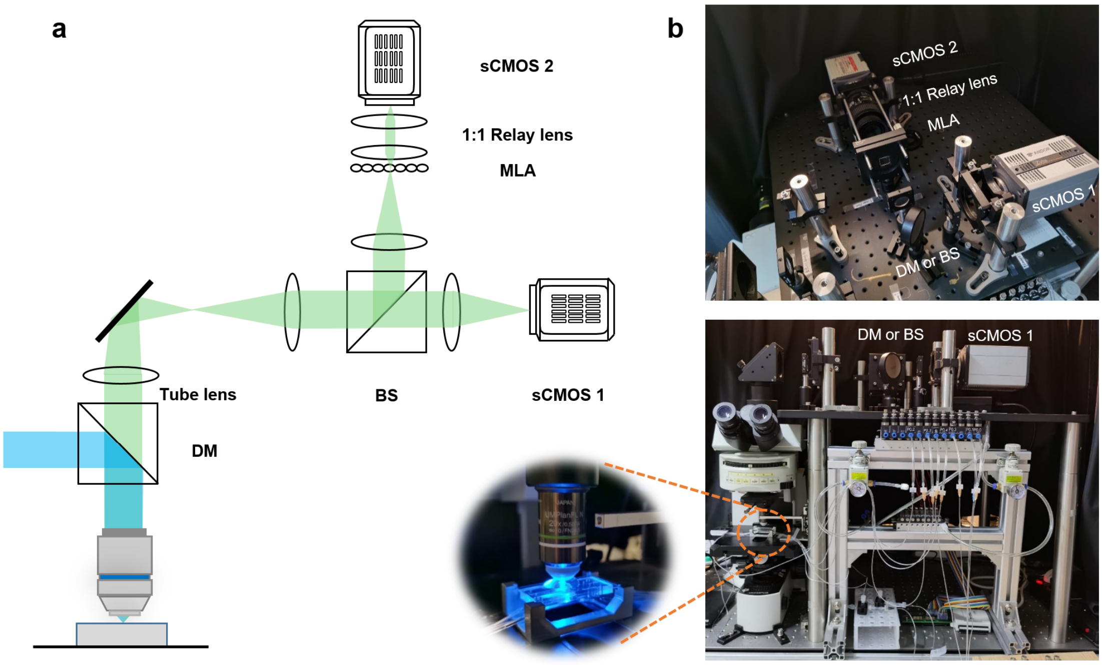


**Fig. S1 Optical setup of the light-field and wide-field microscopy system. (a)** Schematic 2D design of the microscope, showing the optical path with main components. Using a BS (Beam splitter), the wide-field and light-field images can be recorded simultaneously with sCMOS cameras 1 and 2, respectively. A microlens array (MLA, APO-Q-P150-F3.5 (633), OKO Optics) was placed on the plane conjugating to the original image plane of the upright microscope (Olympus, BX51) to generate light field images. The back focal plane of the MLA was imaged with a 1:1 relay system (AF 60 mm 2.8D, Nikon) onto the sensor chip of sCMOS 2. **(b)** Photographs of our imaging system from the top and front views. The magnified photo shows the close-up view of the sample area where a microfluidic chip was used to host the worms within the FOV of 20× objective.

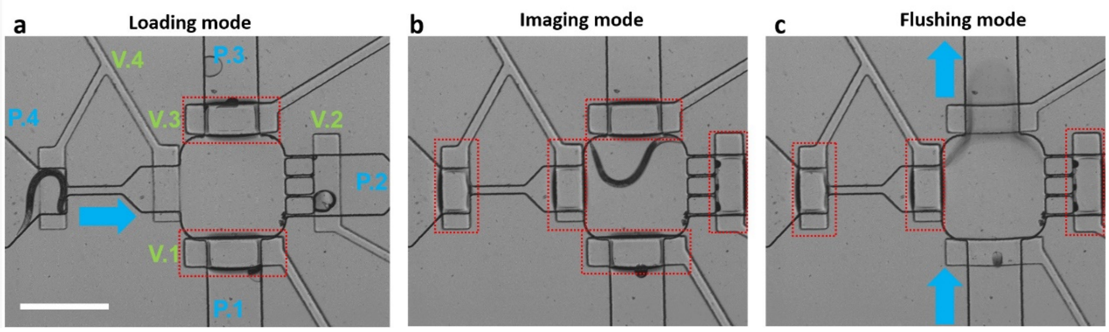


**Fig. S2 Bright-field images of the chip working under loading mode (a), imaging mode (b) and flushing mode (c).** Blue arrows indicate direction of worm's movement. Red dotted boxes indicate the valves that are closed under current mode. Scale bar, 500 μm.

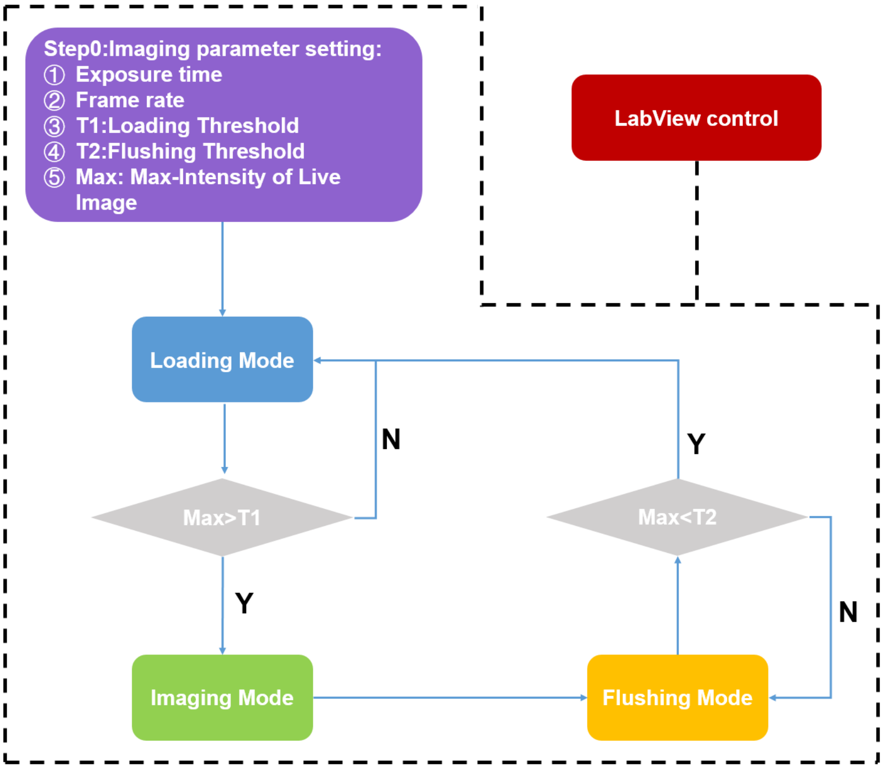


**Fig. S3 Workflow of LabVIEW control program**. First, four parameters, exposure time, frame rate, loading threshold, flushing threshold are set to initialize the control procedure. The loading threshold (T1) and flushing threshold (T2) are two values set lower than the signal intensity of *C. elegans*' and higher than the intensity of background, respectively. Therefore, once a worm is loaded into the imaging chamber with the maximum intensity value of recorded image being larger than the loading threshold (T1), the chip is switched from worm loading mode to imaging mode with corresponding micro valve operation executed. After one-minute observation and the current worm being flushed outside the imaging chamber, the max intensity of the blank background image is turned to be lower than the flushing threshold (T2). The chip is then switched from worm flushing mode to another round of loading mode to push the next worm inside the chamber. Under this control logic, the worm loading, imaging and flushing modes can be cycled automatically.

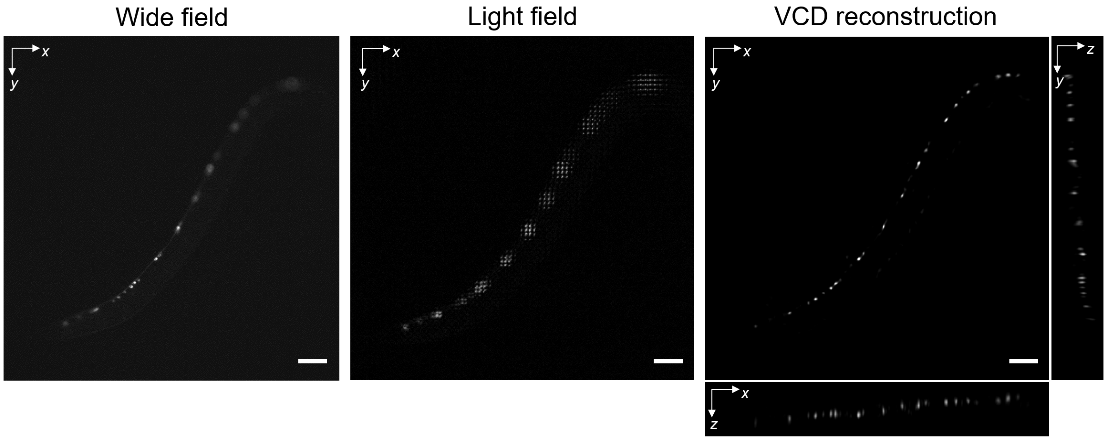


**Fig. S4 Comparison of imaging results of the same moving *C.elegans* by wide field microscopy (left), raw light-field microscopy (LFM, middle) and light-field microscopy combined with VCD reconstruction (VCD-LFM, right)**. Due to the fast locomotion of the worm and the 3D distribution of the neurons, the 2D wide-field image only clearly records a few neurons at 100-Hz acquisition rate. In contrast, after light-field imaging records all the neurons in a 2D frame also at 100-Hz acquisition rate, VCD network further reconstructs the instantaneous distribution (~10 ms) of these neurons in three dimensions. Scale bar, 50 µm.

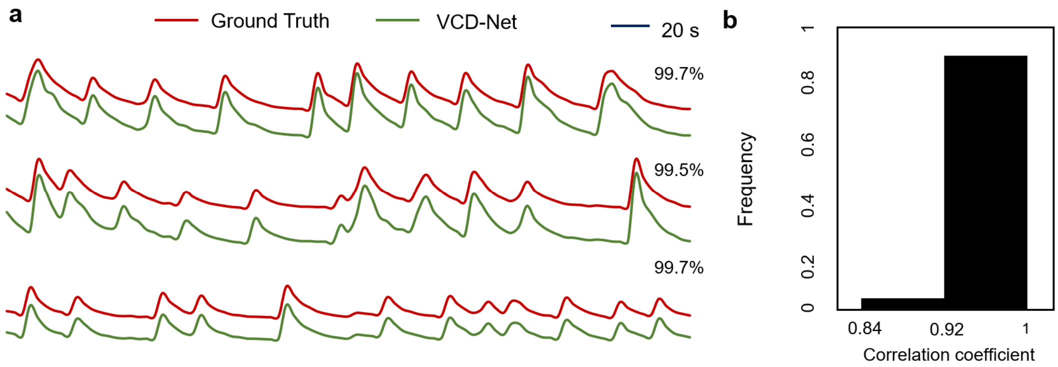


**Fig. S5 Validation of the intensity accuracy of the VCD-reconstructed signals versus ground-truth signals**. We simulated the blinking signals which were also freely moving in 3D space based on the ground-truth status of a GCaMP6-labelled behaving worm. The time-varying signals was then projected into the light-field images and was reconstructed by the trained VCD network. **(a)** Comparison between the signal intensity fluctuations of the ground truths (red lines) and VCD reconstructions (green lines) throughout a 6-minute time period. **(b)** Statistics of the correlation coefficients between the VCD reconstructions and ground truths quantitatively verifies the accuracy of our approach. The chart shows the results of n = 48 simulated signals, with 46 of them above 0.92.

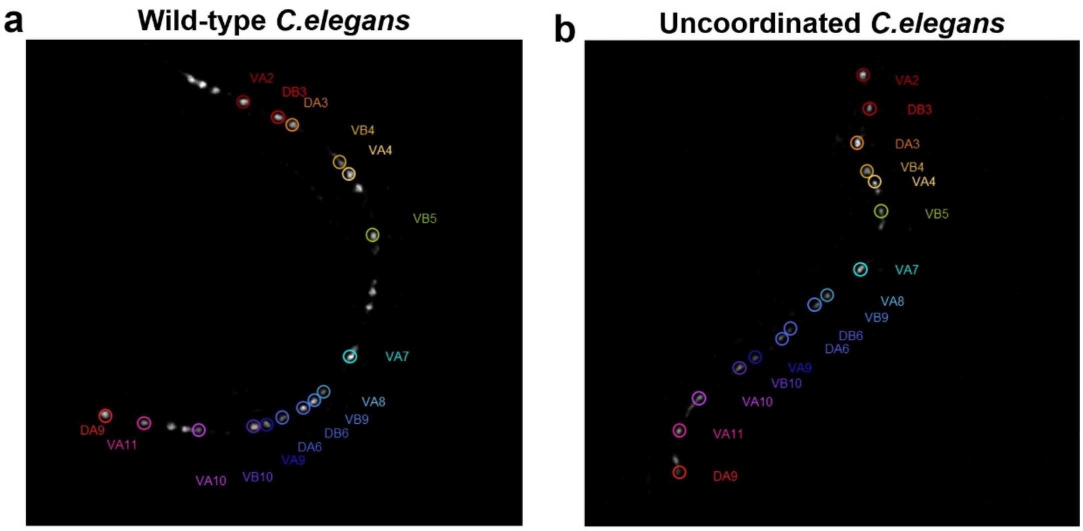


**Fig. S6 Identification of the motor neurons in wild-type (WT) and uncoordinated-type (Unc) worms.** The soma positions of the motor neurons were indicated in one instantaneous maximum-intensity-projection (MIP) of WT and Unc worms, as shown in **(a)** and **(b)**, respectively.

#### Supplementary Videos

**Video S1.** Loading, imaging and flushing procedure of the microfluidic chip.

**Video S2.** Light-field recording of the sequentially loaded worms.

**Video S3.** Raw 2D light-field video of a moving *C. elegans* and its corresponding 3D VCD-reconstruction video.

**Video S4.** Locomotion comparison between a wild-type *C. elegans* and an uncoordinated *C. elegans*.
